# Accounting for cell lineage and sex effects in the identification of cell-specific DNA methylation using a Bayesian model selection algorithm

**DOI:** 10.1101/124826

**Authors:** Nicole M. White, Miles C. Benton, Daniel W. Kennedy, Andrew Fox, Lyn R. Griffiths, Rodney A. Lea, Kerrie L. Mengersen

## Abstract

Cell- and sex-specific differences in DNA methylation are major sources of epigenetic variation in whole blood. Failure to account for these confounders may lead to substantial bias in the identification of differentially methylated CpGs and predicted levels of differential methylation. Previous studies have provided evidence of cell-specific methylation, but all of these have been restricted to the detection of differential methylation in a single cell type. We developed a Bayesian model selection algorithm for the identification of cell-specific methylation profiles that incorporates knowledge of shared cell lineage, to accommodate differential methylation in one or more cell types. Under the proposed methodology, sex-specific differences in methylation by cell type are also assessed. Using publicly available cell-sorted methylation data, we show that 51.3% of female CpG markers and 61.4% of male CpG markers identified were associated with differential methylation in more than one cell type. The impact of cell lineage on differential methylation was also highlighted. An evaluation of sex-specific differences revealed marked differences in CD56^+^NK methylation, within both single and multi-cell dependent methylation patterns. Our findings demonstrate the need to account for cell lineage in studies of differential methylation and associated sex effects.

## Introduction

DNA methylation is a widely studied epigenetic modification that plays an essential role in the regulation of gene expression [1], cell differentiation [2] and the maintenance of chromatin structure [3]. Advances inhigh-throughput technologies [4–6] have made possible the collection of DNA methylation at the genome scale, allowing its relationship with biological processes to be interrogated. Studies of DNA methylation levels in humans, at both the global and individual CpG levels, have revealed associations between aberrant methylation profiles and disease susceptibility [7], including carcinogenesis [8]. These studies have largely been conducted using samples of whole blood, in light of its convenience as a source of DNA and viability as a surrogate for target tissue. Its use, however, presents challenges for analysis and interpretation [9].

The role of DNA methylation in haematopoiesis has motivated the search for cell-specific methylation profiles at the individual CpG level and their association with lineage-specific gene expression [10–12]. To date, the discovery of cell-specific CpG markers has been based on the comparison of methylation levels between select purified cell subtypes [12] or against methylation levels observed in whole blood [13]. In [13], DNA methylation levels for seven purified cell subpopulations were compared and significant differences were identified between Lymphocyte and Myeloid cell populations. The definition of cell-specific methylation in this study and others was restricted the identification of differential methylation in a single cell type. The impact of accounting for more complex methylation profiles based on shared hematopoietic lineage remains unexplored.

Evidence of sex effects in DNA methylation is mixed and studies to date have focused primarily on whole blood. Previous studies of global and autosomal CpG methylation have reported a tendency for higher methylation in males [14] and sex-specific differences at varying numbers of CpG probes, across different chromosomes [15, 16]. A meta-analysis of published findings [17] identified 184 sex-specific, autosomal CpG probes, although average methylation differences were small and included studies did not account for cellular heterogeneity. Evidence for sex-specific methylation patterns for purified cell subtypes remains relatively unexplored. A recent study [18] examined sex-specific methylation levels for four purified immune cell subtypes (B cells, Monocytes, CD4+Foxp3−,CD8+T), and their role in immune-mediated diseases. Cell-specific methylation patterns in females versus males were compared and sex-specific differences were identified for each cell type. Similar to previous studies of cell-specific methylation, the identification of these differences was restricted to a single cell type, therefore precluding the exploration of sex-specific differences as a function of cell lineage.

In light of current limitations, this paper proposes new methodology for the identification of cell-specific methylation profiles at the individual CpG level and associated sex effects. Our methodology incorporates knowledge about cell lineage to accommodate the identification of differential methylation in one or multiple cell types, therefore expanding upon the range of methylation patterns currently studied. The main component of the methodology is a model choice algorithm, formulated within a Bayesian framework, which allows evidence for a range of cell-specific models to be assessed simultaneously. Using cell-sorted methylation data from female and male samples, panels of cell-specific CpG markers are identified for each sex and common marker panels are derived. Subsequent analysis of identified CpG markers demonstrate the need to account for lineage in the discovery of cell-specific methylation pattern.

## Materials and methods

### Data

Publicly available Illumina 450K methylation data were obtained from six healthy males subjects [13]. The data contained methylation *β*-values on seven isolated cell populations: CD19B^+^ cells, CD14^+^ Monocytes, CD4^+^T cells, CD8^+^T cells, CD56^+^ Natural Killers (NK), Neutrophils (Neu) and Eosinophils (Eos). Samples of the same cell types excluding Eosinophils were also obtained for five healthy female subjects [18]. In the absence of Eosinophils, The methodology described in this paper was therefore only applied to the six cell types common to both datasets. For both sexes, 445,603 CpG probes were available for analysis.

Consistent with previous findings [13], differences in DNA methylation levels across CpG sites were primarily driven by cell type differences, as opposed to being an artefact of between-subject variation (Fig 1). A hierarchical clustering of cell-sorted samples revealed that differences in methylation levels were closely aligned with expected cell lineage in whole blood and were sex-independent.

**Figure 1:**
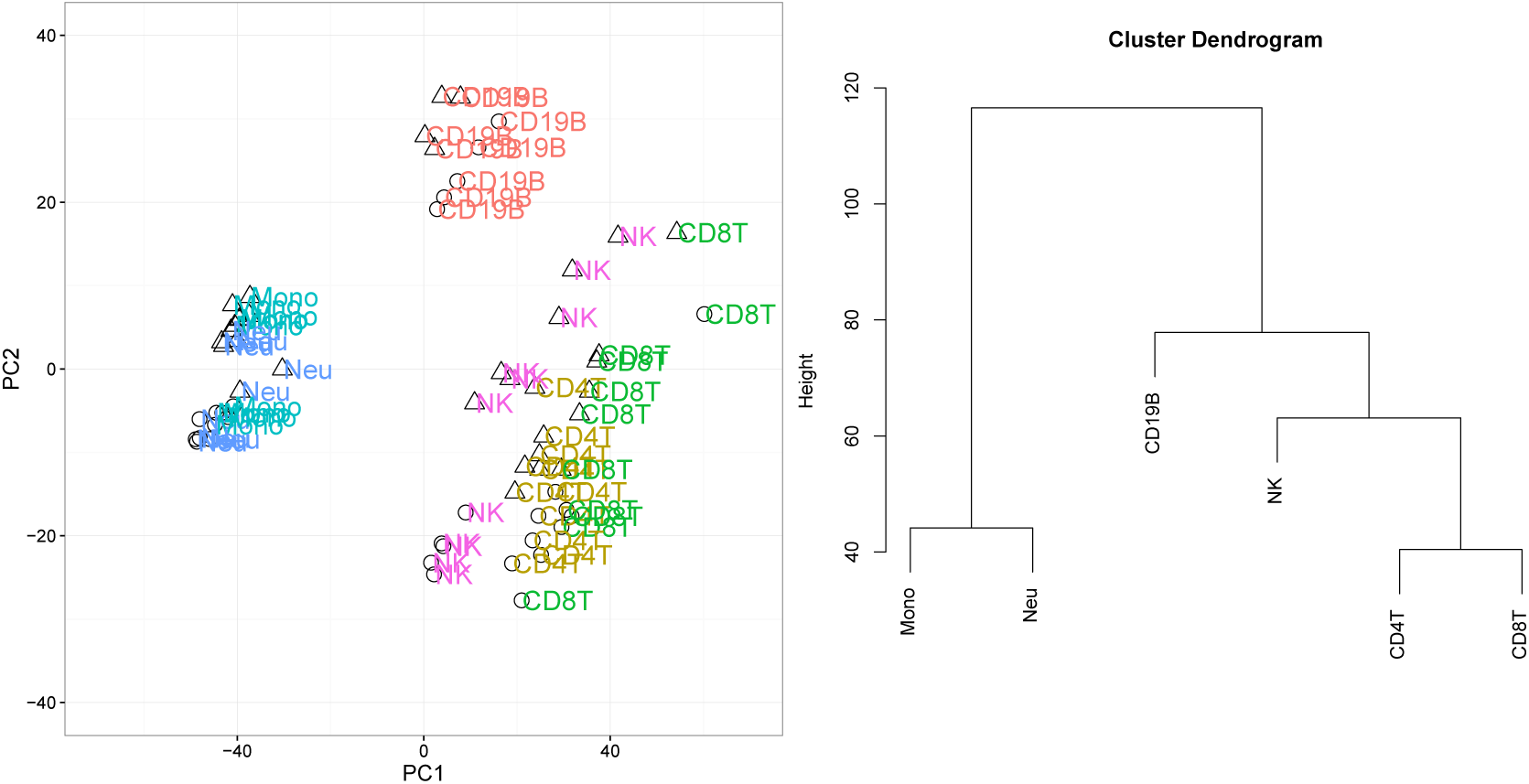
Left: Principal components plot over samples (Circle = Female, Triangle = Male). Right: Dendrogram of cell-sorted DNA methylation samples over 445,603 CpG probes for female and male samples. The response variable was equal to the mean methylation *β*–value by each cell type, at each CpG probe.

Additional cell sorted 450K methylation data on female and male subjects [18], aged 32-50 years, were obtained from public databases, for validation of model results. Female data consisted of cell-sorted samples on six healthy subjects downloaded from ArrayExpress (Accession number: E-ERAD-179) for CD19B^+^B cells, CD14^+^ Monocytes, CD4^+^T and CD8^+^ T cells. Methylation data on six healthy male subjects from the same study were downloaded from Gene Expression Ominibus (Accession number: GSE71245). Following filtering and normalisation, 436,067 and 445,307 CpG sites were available for validation for females and males respectively, based on their correspondence with available sites in the training data.

### Model Formulation

The methodology presented in this paper was motivated by the identification of cell-specific methylation at the individual CpG level. A cell-specific CpG was defined as any CpG probe where the expected methylation level was different for one (single cell-specific) or more (multi- cell-specific) cell types, relative to others observed. The true cell-specific methylation pattern at each CpG probe was assumed unknown and treated as a model selection problem, with key details outlined in this Section.

Information about cell lineage inferred from available samples was used to define candidate models for selection at each CpG. Using the results of hierarchical clustering (Fig 1), eleven candidate models were defined (Table 1) where each model represented a dendrogram split or node. Each candidate model therefore proposed a different cell-specific methylation pattern, characterised by a unique partition of cell types into non-overlapping groups. For a given candidate model, cell types assigned to the same group were assumed to share the same level of methylation. Two additional models, corresponding to the null and saturated cases, were also considered.

**Table 1:**
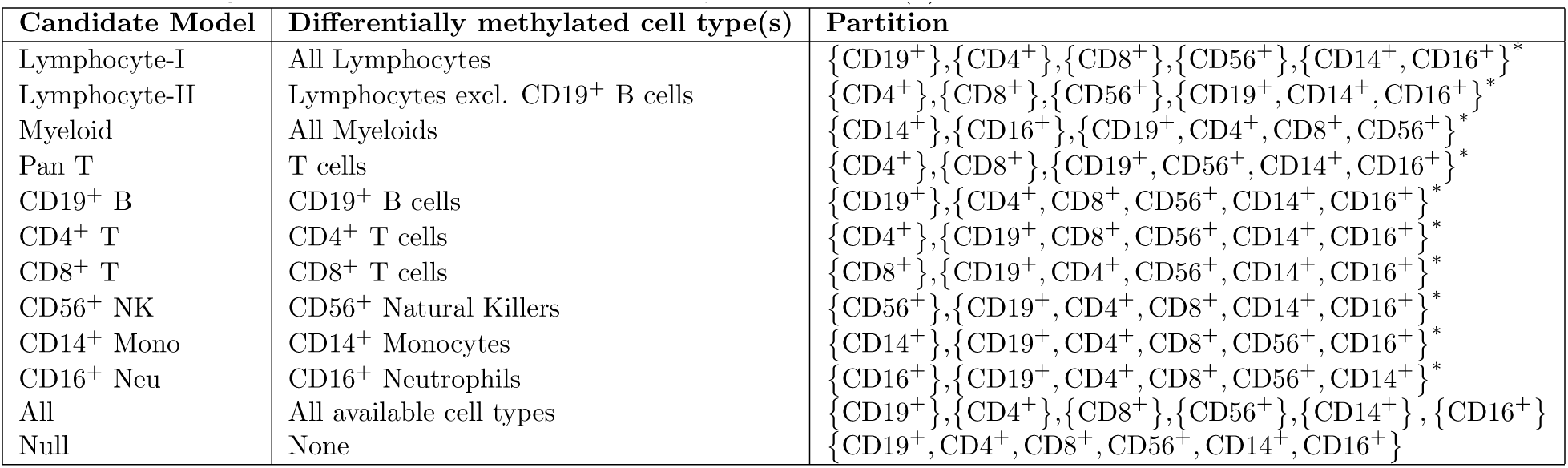
Description of cell-specific models based on cell lineage from Fig 1. Each candidate model corresponds to a partition of cell types into non-overlapping groups. Cell types within each set of parentheses, {}, belong to the same partition and were assumed to have the same level of methylation. For each cell-specific model excluding ‘All’, the partition annotated by an asterisk (*) denotes the reference partition.

Each candidate model was translated into a linear model, where each cell type grouping was represented by a different mean parameter. For sex *s*, observed methylation *β*–values for each cell type at CpG *k* were represented by the vector, y_*iks*_ = (*y*_*iks1*_,…, *y*_*iksj*_), for samples *i* = 1,…, *n*_*s*_. The partition defined under each candidate model was encoded into the design matrix, **X**_*m*_, of size *J* × *P*^(*m*)^, where *P*^(*m*)^ was equal to the number of partitions for candidate model m. Mean methylation levels for each partition were represented by the vector 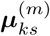 of length *P*^(*m*)^. The likelihood for a single CpG probe *k* = *1*,…, *K* under candidate model m was given by,

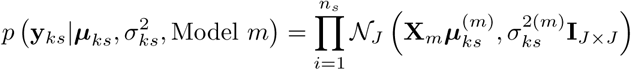
where 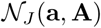 defines a *J*–dimensional Normal distribution with mean vector a and variance-covariance matrix **A**. The unknown variance was assumed to be common across all cell types, denoted by 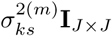 where **I** is the identity matrix. This variance-covariance structure was chosen for identifiability reasons given sample sizes available.

A Bayesian approach to model selection was adopted, which allowed for probabilistic statements to be made about the relative fit of each candidate model. Under this approach, the posterior probability of each model conditional on the observed data was calculated for all candidate models. The posterior probability of model *m* compared to other candidates *m*′ was calculated as,

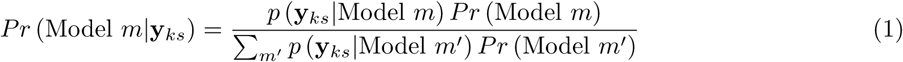
where the sum of probabilities over all candidate models was equal to 1. The term *p* (y_*ks*_|Model *m*) was obtained by integrating over all unknown parameters from the likelihood and prior distributions. Prior probabilities of each model, *Pr* (Model *m*), were assumed to be equal to reflect *a* lack of model preference *a priori*.

Prior distributions for remaining parameters were selected such that Eq 1 could be derived analytically. A g-prior distribution [19] was adopted for each 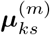,

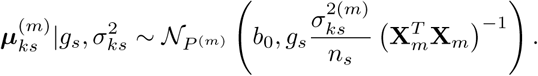

The g-prior is a popular choice in linear model selection settings, as it allows the experimenter to introduce information on the scale of **X**_*m*_. Expected methylation levels were centered around the overall methylation level, *b_0_*, with variance proportional the standard error of each partition. The scaling factor *g_s_* > 0 is interpreted as a relative weighting of the prior versus the observed data. Here, *g_s_* was assumed common over all CpG probes and estimated by maximising the marginal likelihood averaged over candidate models. In this paper, *b_0_* was set to the global methylation mean, averaged over CpGs and cell types. A conjugate prior distribution for 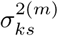 of form 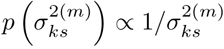 completed model specification [19].

The expectation-maximisation (EM) algorithm was used to obtain a global Empirical Bayes estimate for *g_s_* [20]. This computational approach provided significant computational benefits over sampling-based approaches, namely Markov chain Monte Carlo (MCMC), and was possible given closed-form solutions for *P* (y_*ks*_|Model *m*) conditional on *g_s_*. In addition, the desired posterior model probabilities from Eq 1 were available upon convergence of the EM algorithm. The proposed methodology was implemented in R, with code available as Supporting Information (File S1).

### Making the most of the Bayesian approach for model based inference

The accommodation of model and parameter uncertainty under the Bayesian approach formed the basis of subsequent inference, namely the identification of cell-specific CpGs, the estimation of differential methylation by cell type and the assessment of sex-specific differences by cell type. Brief details of each inference are provided in this Section.

#### CpG marker identification

CpG probes or ‘markers’ associated with each cell-specific pattern were identified by comparing posterior model probabilities at each CpG probe. For each candidate model and sex, markers were identified by reviewing the set of *K* model probabilities: *Pr* (Model *m*|y_1*s*_),…,*Pr* (Model *m*|y_*Ks*_). A 5% Bayes’ False Discovery Rate (FDR) [21] was applied to each set of probabilities to control the expected number of false discoveries. Common CpG markers were defined as CpGs that were identified in both female and male samples, for the same candidate model. The compilation of common marker panels allowed for the assessment of sex-specific differences by cell type, within each candidate model.

#### Estimation of differential methylation

Posterior inference about each marker panel focused on the estimation of differential methylation by cell type relative to its corresponding reference partition (Table 1). Under model *m*, the posterior distribution of differential methylation for cell-specific partition *p* and sex *s* is,

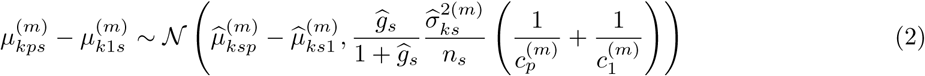
where *c_p_* is the number of cell types in partition *p* and 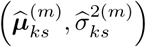 are posterior estimates of each mean and residual variance, respectively, for which direct estimates were available. For CpGs identified under each marker panel, Eq 2 was summarised in terms of a posterior mean and 95% credible interval (CI). The posterior probability of differential methylation being within selected ranges (<0.10, 0.10-0.20,0.20-0.30,0.30,0.40,0.40-0.50,>0.5), was also calculated.

The posterior distribution in Eq 2 was also used to validate selected CpG markers. Validation of common CpG markers was limited to CD19^+^ B, CD4^+^ T, CD8^+^ T and Pan T models, in light of validation samples available. Using all validation samples for each sex, the average *β*–value for each cell type and CpG was computed. The difference between each average *β*–value and reference partition was then compared with the corresponding 95% CI from Eq 2 for the appropriate candidate model. Concordance rates with respect to the predicted direction of methylation (hypomethylated, hypermethylated) for validation samples were also calculated. This joint approach was motivated by the limitation that validation based on coverage of 95% CIs relied on a posterior estimate of 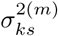. When this estimate is small and/or underestimated, proportions of validated markers based on CI coverage only were likely to be low, even if differences in methylation between training and validation samples were small. Finally, it is noted that not all cell types observed in the training data were available in the validation samples. Given the properties of the multivariate Normal distribution, this discrepancy was addressed by using the appropriate marginal distributions for the available cell types.

#### Evaluation of sex effects

A similar expression to Eq 2 was derived to evaluate sex-specific methylation differences within common CpG marker panels. In this case, focus was on the comparison of female and male methylation estimates for differentially methylated cell types, as defined by the corresponding candidate model. The posterior distribution of this difference across model partitions was Multivariate Normal,

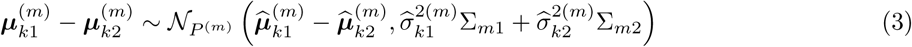
where 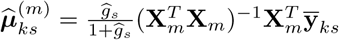 for *s* = 1, 2. To assess evidence for sex effects, the posterior probability that the difference in methylation between sexes using Eq 3 was at least 0.10 was calculated for each cell-specific partition. This calculation again relied on estimates of the residual variance, 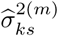 for each sex which were set to their posterior mean estimate. A sex-specific difference for a given partition was declared if the posterior probability of a difference greater than 0.10 exceeded 0.95.

### CpG markers to genes: Assessment of SNP effects, genomic features and pathway enrichment

To provide additional evidence to support our method of marker identification, a pathways enrichment analysis was performed to explore the underlying biology of gene sets derived from common CpG marker panels. Common CpG markers were mapped to genes using Illumina Human Methylation 450k annotation data available from Bioconductor [22]. SNP information was also collated to infer the percentage of SNP associated markers with associations limited to SNPs located directly on the CpG loci. Using KEGG functional analysis in WebGestalt [23], a hypergeometric test was applied to each marker panel. A 5% FDR [24] was applied to resulting p-values to identify significant pathway enrichment for derived gene lists.

## Results

### Cell lineage impacts the identification of differentially methylated CpGs

Across all cell-specific models, 83,449 and 97,747 CpG markers were identified for females and males, respectively (Table 2). Among female samples, 42,834 markers (51.3%) were associated with differential methylation in multiple cell types, which included a three-fold increase in Pan T markers compared with males. A larger proportion of multi- cell dependent markers among males was observed (64.8%), due to larger numbers identified under Lymphocyte-II and Myeloid models. Among single cell dependent models, CD19^+^B was the most frequently observed marker type for both sexes, with 25,611 in females and 18,271 in males.

**Table 2:** Number of CpG markers identified by cell dependent model for females versus males, based on the applications of a 5% Bayes False Discovery Rate (FDR). For each model, the number of common markers identified in both sexes is also listed. The number of CpGs identified for a single sex only are given in brackets for each marker panel.

| Candidate model | Female | Male | Common |
| --- | --- | --- | --- |
| All | 3873 (1699) | 3553 (1379) | 2174 |
| Myeloid | 12541 (3637) | 29534 (20630) | 8904 |
| Lymphocyte-I | 15596 (7369) | 14105 (5878) | 8227 |
| Lymphocyte-II | 4241 (1032) | 14015 (10806) | 3209 |
| Pan T | 6583 (5546) | 2126 (1089) | 1037 |
| CD19<sup>+</sup> B | 25611 (11987) | 18271 (4647) | 13624 |
| CD4<sup>+</sup> T | 1277 (821) | 2050 (1594) | 456 |
| CD8<sup>+</sup> T | 2299 (1985) | 2742 (2428) | 314 |
| CD56<sup>+</sup> NK | 4893 (3302) | 4346 (2755) | 1591 |
| CD14<sup>+</sup> Mono | 1383 (627) | 1681 (925) | 756 |
| CD16<sup>+</sup> Neu | 5152 (2992) | 5324 (3164) | 2160 |

A total of 42,452 CpGs were identified under the same cell-specific methylation pattern for both sexes, corresponding to 9.5% of the observed methylome. Within this subset, 23,551 (55.5%) were defined by differential methylation in more than one cell type. Over all candidate models, smaller frequencies of common markers were associated with T-cell dependent markers (CD4^+^, CD8^+^, Pan T).

### Differential methylation is affected by cell lineage among common CpG markers

Common CpG markers associated with differential methylation in Myeloid cell types (CD14^+^ Mono, CD16^+^ Neu) were consistently hypomethylated across all relevant marker panels (Table 3). Among Lymphocytes, CD8^+^T markers were the least likely to be hypomethylated (35.99%). Smaller proportions of hypomethylation among Lymphocyte I and II panels were indicative of greater methylation among Lymphocyte versus Myeloid cell subtypes.

**Table 3:** Percentage of hypomethylated common markers by cell type within single cell-specific (CD19^+^B, CD4^+^T, CD8^+^T, CD56^+^NK) and multi- cell-specific (Myeloid, Lymphocyte-I, Lymphocyte-II, Pan T) marker panels.

|  | Sex | Single cell-specific markers |  |  |  |  |  |
| --- | --- | --- | --- | --- | --- | --- | --- |
|  |  | CD19<sup>+</sup>B | CD4<sup>+</sup>T | CD8<sup>+</sup>T | CD56<sup>+</sup>NK | CD14<sup>+</sup>Mono | CD16<sup>+</sup>Neu |
|  | Female | 58.24 | 65.57 | 35.99 | 90.19 | 94.71 | 96.39 |
|  | Male | 58.24 | 65.57 | 35.99 | 90.19 | 94.71 | 96.39 |
|  | Sex | Multi- cell-specific markers |  |  |  |  |  |
|  |  | CD19<sup>+</sup>B | CD4<sup>+</sup>T | CD8<sup>+</sup>T | CD56<sup>+</sup>NK | CD14<sup>+</sup>Mono | CD16<sup>+</sup>Neu |
| Myeloid | Female | – | – | – | – | 93.53 | 93.41 |
|  | Male | – | – | – | – | 93.53 | 93.41 |
| Pan T | Female | – | 43.30 | 43.30 | – | – | – |
|  | Male | – | 43.30 | 43.30 | – | – | – |
| Lymphocyte-I | Female | 29.52 | 26.70 | 27.26 | 29.11 | – | – |
|  | Male | 29.52 | 26.70 | 27.46 | 28.86 | – | – |
| Lymphocyte-II | Female | – | 44.13 | 45.56 | 45.72 | – | – |
|  | Male | – | 44.13 | 45.65 | 45.81 | – | – |

The impact of cell lineage on differential methylation was greatest among marker panels related to Lymphocytes (Fig 2). Lower levels of differential methylation ( <0.10) were concentrated within single cell dependent markers (CD19^+^B, CD4^+^T, CD8^+^T, CD56^+^NK). In contrast, the comparison of distributions indicated a wider range of posterior estimates observed for Pan T, Lymphocyte-I and Lymphocyte-II panels. The comparison of distributions associated with CD14^+^ Monocytes showed little evidence of being affected by cell lineage.

**Figure 2:**
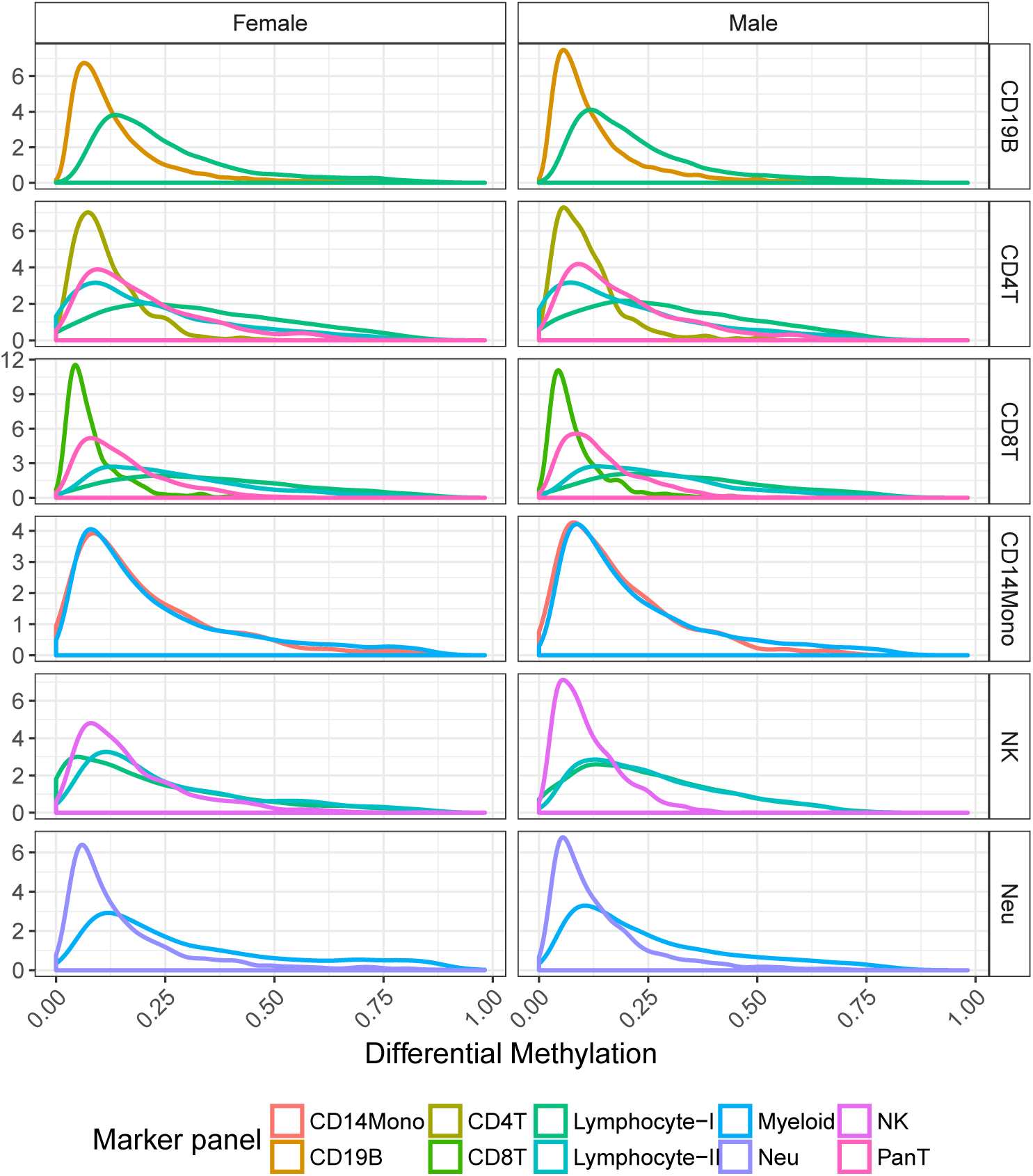
Distribution of posterior mean estimates of differential methylation for each purified cell type across corresponding marker panels. Posterior estimates are summarised for females (first column) and males (second column) across common CpG markers.

The distribution of differential methylation by sex among common Pan T markers revealed considerable variation for both hypermethylated and hypomethylated states compared to CD4^+^T and CD8^+^ T panels (Fig 3). Differential methylation levels among Pan T markers tended to be greater for CD4^+^ T cells; 25 markers in this subset showed strong evidence of differential methylation greater than 0.5 for both sexes (Fig S1).

**Figure 3:**
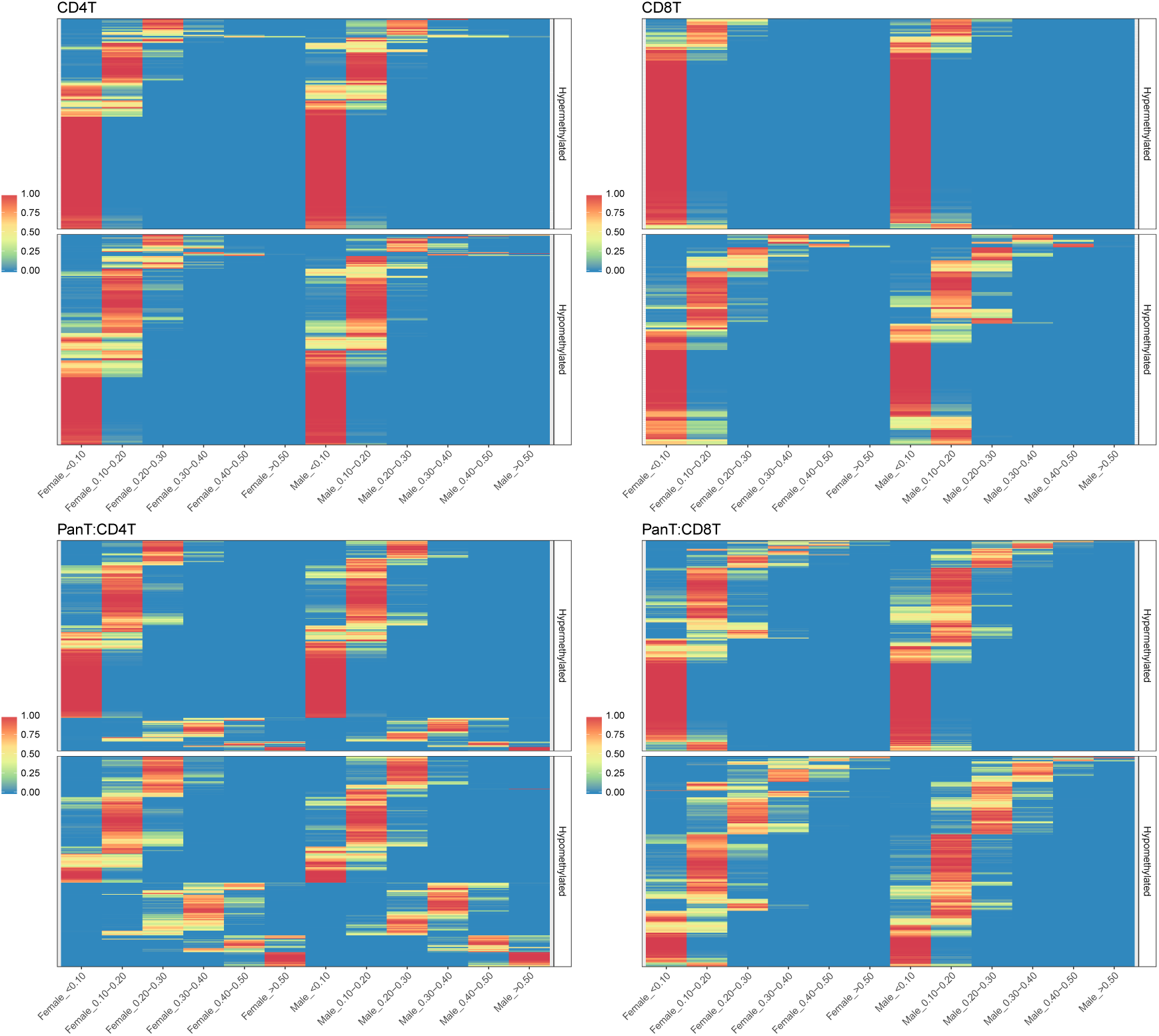
Posterior probability heatmaps for varying levels of differential methylation for common CD4^+^T, CD8^+^T and Pan-T markers. For each cell type, CpG markers (rows) are ordered the same for males and females, to enable sex-specific comparisons. Markers are further classified by methylation state (Hypomethylated, Hypermethylated), based on their posterior mean estimate of differential methylation. First row (L-R): CD4^T^, CD8^+^T; Second row: Pan T.

### Validation of common CpG markers for B- and T-lymphocytes

The validation of common markers with respect to mean differences was generally higher in males than females, across all immune cell subtypes (Table 4). Whilst validation based on coverage of credible intervals was low, differences between training and validation estimates were relatively small across all markers tested; for CD4^+^T markers, approximately 80% of absolute differences were less than 10% (Fig S2). Furthermore, comparisons with respect to inferred methylation state showed very high concordance rates for all marker panels and these findings were consistent for both sexes.

**Table 4:**
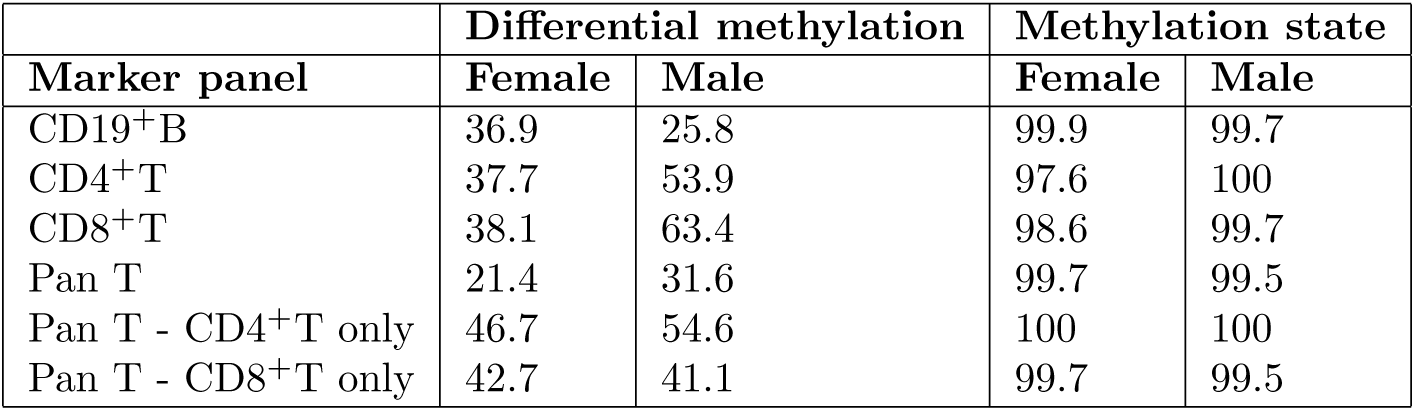
Percentage of common CpG markers that were classified as validated based on the coverage of 95% credible region inferred from the training data, by sex and marker type. The outcome of interest for validation was estimated level of differential methylation. For Pan T markers, results are summarised by individual cell type and the jointly, the latter corresponding to both CD4^+^T and CD8^+^T differences being validated for the same marker.

| Marker panel | Differential methylation |  | Methylation state |  |
| --- | --- | --- | --- | --- |
|  | Female | Male | Female | Male |
| CD19<sup>+</sup>B | 36.9 | 25.8 | 99.9 | 99.7 |
| CD4<sup>+</sup>T | 37.7 | 53.9 | 97.6 | 100 |
| CD8<sup>+</sup>T | 38.1 | 63.4 | 98.6 | 99.7 |
| Pan T | 21.4 | 31.6 | 99.7 | 99.5 |
| Pan T - CD4<sup>+</sup>T only | 46.7 | 54.6 | 100 | 100 |
| Pan T - CD8<sup>+</sup>T only | 42.7 | 41.1 | 99.7 | 99.5 |

### Genomic feature distributions and enrichment analysis

The effects of cell lineage were evident in the comparison of genomic features, with higher proportions of markers residing in Transcription Start Sites (TSS) for Lymphocyte cell subtypes compared with Myeloids (Table 5). Common markers were concentrated in the gene body and intergenic regions, with 54.73% of CD56^+^NK markers located in the gene body. Among Lymphocyte cell subtypes, TSS proportions were highest for T cells, with 18.28% and 16.48% for CD4^+^T and CD8^+^ T cells, respectively.

**Table 5:** Distribution of genomic features among common CpG markers by cell-specific methylation pattern.

| Marker panel | 1stExon | 3'UTR | 5'UTR | Body | Intergenic | TSS1500 | TSS200 |
| --- | --- | --- | --- | --- | --- | --- | --- |
| Myeloid | 2.88 | 5.90 | 11.17 | 46.85 | 15.41 | 13.65 | 4.14 |
| Lymphocyte-I | 4.10 | 4.20 | 14.26 | 40.84 | 13.26 | 17.05 | 6.28 |
| Lymphocyte-II | 3.58 | 4.31 | 14.48 | 40.68 | 12.92 | 18.15 | 5.88 |
| PanT | 4.32 | 2.19 | 14.29 | 40.75 | 13.26 | 17.98 | 7.20 |
| CD19<sup>+</sup>B | 7.07 | 4.32 | 13.55 | 38.71 | 13.81 | 14.20 | 8.35 |
| CD4<sup>+</sup>T | 3.79 | 3.52 | 15.14 | 41.64 | 12.40 | 18.28 | 5.22 |
| CD8<sup>+</sup>T | 6.67 | 2.78 | 12.59 | 38.52 | 13.15 | 16.48 | 9.81 |
| CD56<sup>+</sup>NK | 2.21 | 5.07 | 12.17 | 54.73 | 12.43 | 9.80 | 3.59 |
| CD14<sup>+</sup>Mono | 1.72 | 4.38 | 10.47 | 47.30 | 20.60 | 11.33 | 4.21 |
| CD16<sup>+</sup>Neu | 1.72 | 8.12 | 10.63 | 53.68 | 12.21 | 10.89 | 2.75 |

Moderate proportions of markers were directly associated with SNPs and these levels were maintained between sex-specific and common marker panels, averaging 30.7% in common markers (Table S1). Higher SNP proportions were observed in marker panels related to the lineage of Myeloid cells, except for CD56^+^NK markers of which 36.02% were SNP associated. For CpG probes not assigned to any common marker panel, a similar degree of SNP association was observed (26.61%).

Significant pathways enrichment for common markers were associated with immune cell subtypes (Table 6) all including biologically relevant pathways.

**Table 6:**
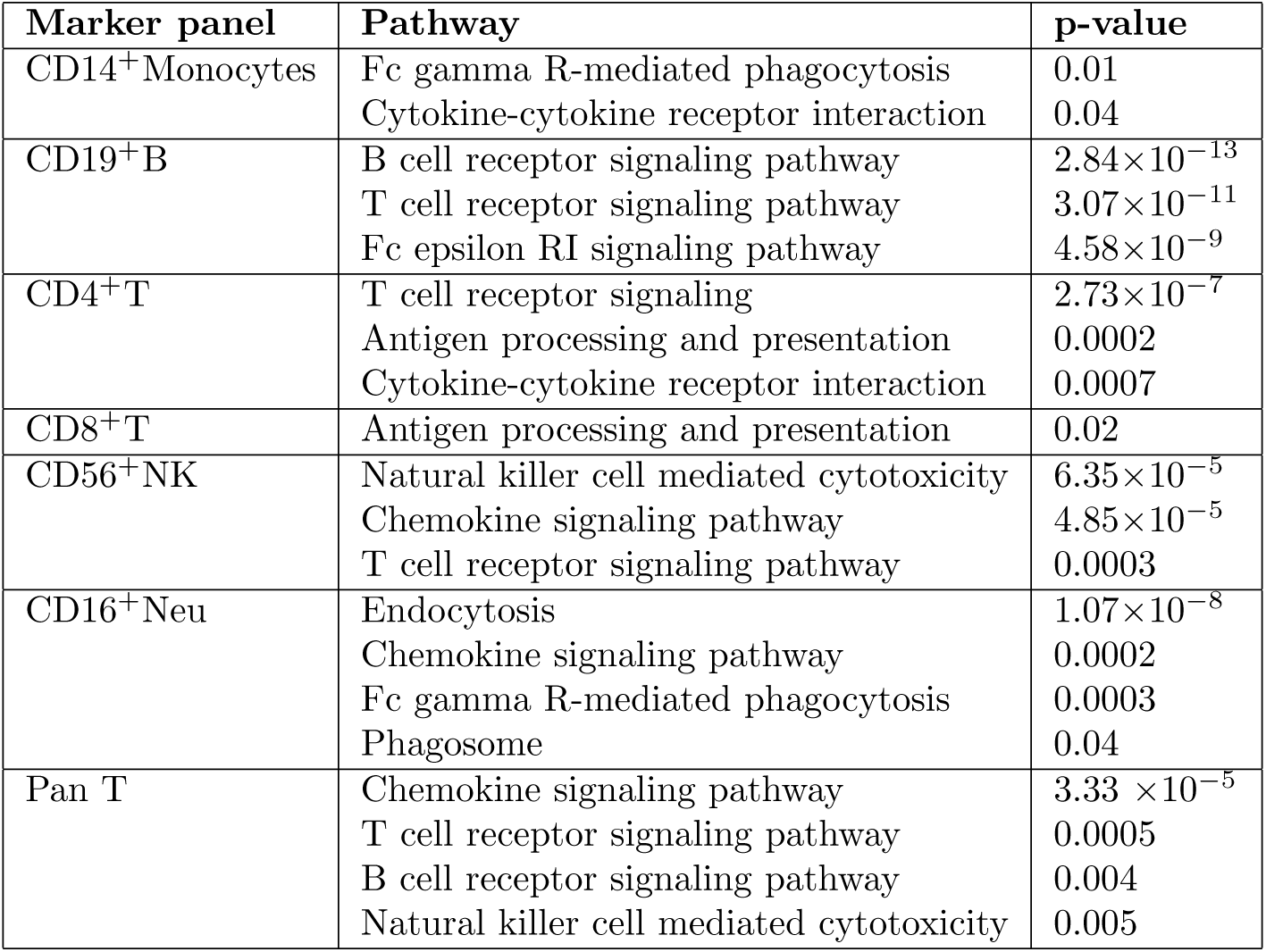
Summary of biologically relevant pathways identified in enrichment analysis of common marker panels. All pathways were identified based on a Benjamini-Hochberg (BH) adjusted p-value < 0.05.

| Marker panel | Pathway | p-value |
| --- | --- | --- |
| CD14<sup>+</sup>Monocytes | Fc gamma R-mediated phagocytosis | 0.01 |
|  | Cytokine-cytokine receptor interaction | 0.04 |
| CD19<sup>+</sup>B | B cell receptor signaling pathway | $2.84 \times 10^{-13}$ |
| | T cell receptor signaling pathway | $3.07 \times 10^{-11}$ |
| | Fc epsilon RI signaling pathway | $4.58 \times 10^{-9}$ |
| CD4<sup>+</sup>T | T cell receptor signaling | $2.73 \times 10^{-7}$ |
|  | Antigen processing and presentation | 0.0002 |
|  | Cytokine-cytokine receptor interaction | 0.0007 |
| CD8<sup>+</sup>T | Antigen processing and presentation | 0.02 |
| CD56<sup>+</sup>NK | Natural killer cell mediated cytotoxicity | $6.35 \times 10^{-5}$ |
| | Chemokine signaling pathway | $4.85 \times 10^{-5}$ |
|  | T cell receptor signaling pathway | 0.0003 |
| CD16<sup>+</sup>Neu | Endocytosis | $1.07 \times 10^{-8}$ |
|  | Chemokine signaling pathway | 0.0002 |
|  | Fc gamma R-mediated phagocytosis | 0.0003 |
|  | Phagosome | 0.04 |
| Pan T | Chemokine signaling pathway | $3.33 \times 10^{-5}$ |
|  | T cell receptor signaling pathway | 0.0005 |
|  | B cell receptor signaling pathway | 0.004 |
|  | Natural killer cell mediated cytotoxicity | 0.005 |

### Assessment of common marker profiles shows sex effects in CD56^+^NK, CD16^+^Neutrophils

The majority of common CpG markers did not exhibit sex-specific differences in methylation profile (Table 7).Differences within CD56^+^NK and Lymphocyte-I/II panels indicated 12.32, 14.15 and 20.64% of respective markers exhibited sex effects, with the majority corresponding to autosomal CpGs. No sex-specific differences were identified among common CD8^+^ T markers.

**Table 7:** Summary of sex-specific differences by common CpG marker panel. A difference between male and female methylation estimates ≥0.10 was the outcome of interest. A marker was declared sex-specific if the posterior probability for this outcome exceeded 0.95, for one or more model-based partitions. For each marker panel, total numbers of sex-specific markers and autosomal sex-specific markers are given.

| Marker panel | Non sex-specific | Sex-specific differences |  |
| --- | --- | --- | --- |
|  | (%) | Total | Total Autosomal |
| Myeloid | 91.69 | 740 | 685 |
| Lymphocyte-I | 79.36 | 1698 | 1635 |
| Lymphocyte-II | 85.85 | 454 | 444 |
| Pan T | 99.42 | 6 | 2 |
| CD19<sup>+</sup> B | 99.75 | 34 | 11 |
| CD4<sup>+</sup> T | 98.90 | 5 | 1 |
| CD8<sup>+</sup> T | 100 | 0 | 0 |
| CD56<sup>+</sup> NK | 87.68 | 196 | 194 |
| CD14<sup>+</sup> Mono | 99.07 | 7 | 4 |
| CD16<sup>+</sup> Neu | 96.67 | 72 | 66 |

Sex-specific differences identified among single cell-specific markers were uniquely mapped to 215 genes (Table S2). The majority of identified cases were concentrated in the CD56^+^ NK common marker panel. Four CD4^+^ T markers were associated with greater methylation observed in females and were mapped to a single gene, *CD40LG*, located on the X chromosome. Annotation information for the full list of sex-specific markers identified is provided in the Supplementary Material (File S2).

sex effects in CD56^+^ NK methylation was prominent in Lymphocyte-I and Lymphocyte-II common marker panels, in addition to the CD56^+^ NK panel (Fig 5). Within the Lymphocyte-I panel, 1698 common CpG markers were identified as sex-specific, of which 1627 showed differences in CD56^+^ NK methylation only. Similarly, 442 common Lymphocyte-II markers showed differences in CD56^+^ NK methylation between sexes. The tendency for CD56^+^ NK methylation to be higher in males was common across all three panels.

**Figure 4:**
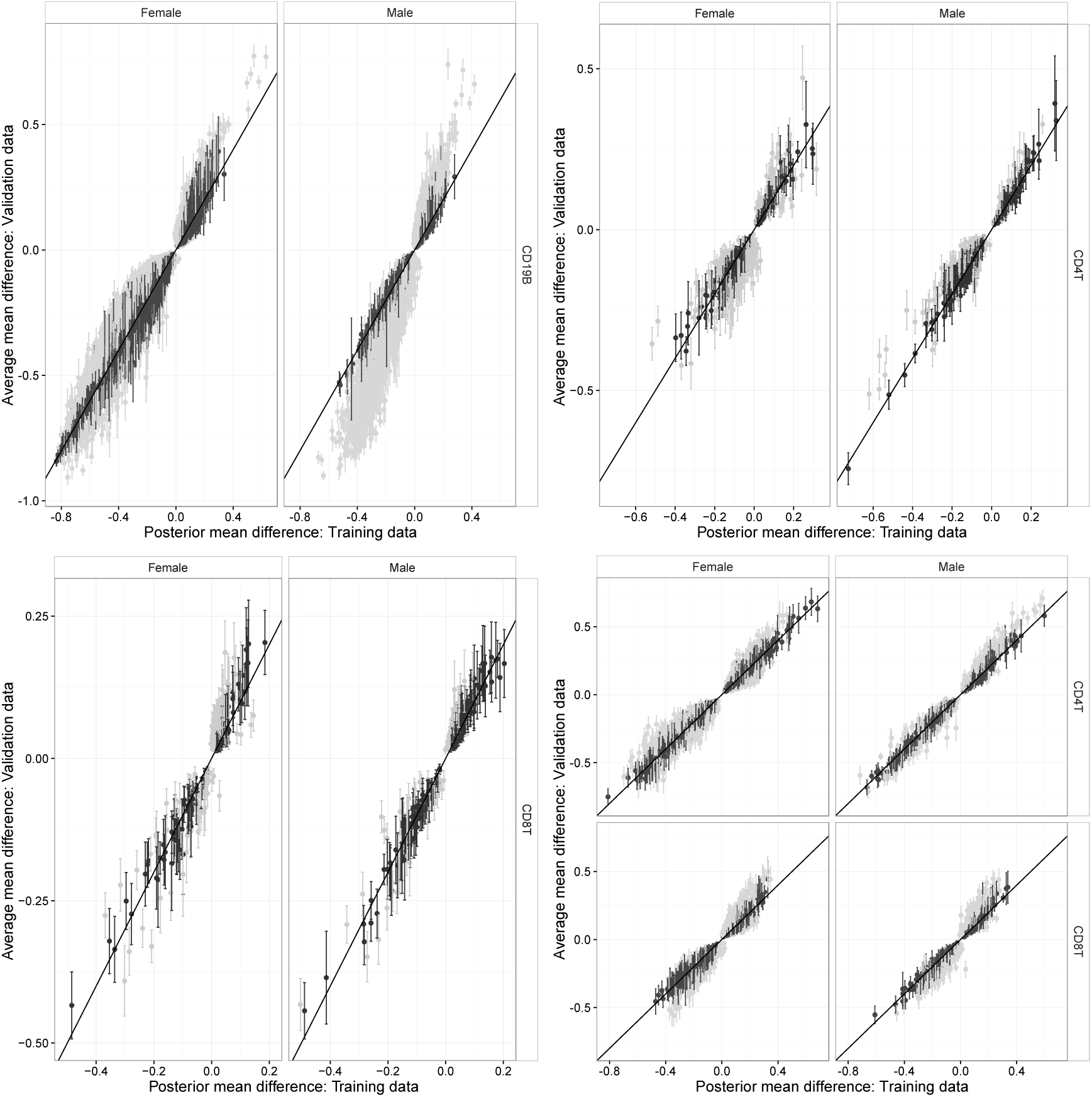
Validation of common marker panels for immune cell subtypes, based on the comparison of mean differences for each cell type relative to its corresponding reference partition. For each marker type, posterior mean differences inferred from the training data (x-axis) are compared with the average mean difference calculated from the validation data (y-axis). Estimates of each posterior mean difference are accompanied by a respective 95% CI. Validated markers based on 95%CI covered are coloured black. First row (L-R): CD19^+^, CD4^+^ T; Second row (L-R): CD8^+^ T, Pan T. CI: Credible interval.

**Figure 5:**
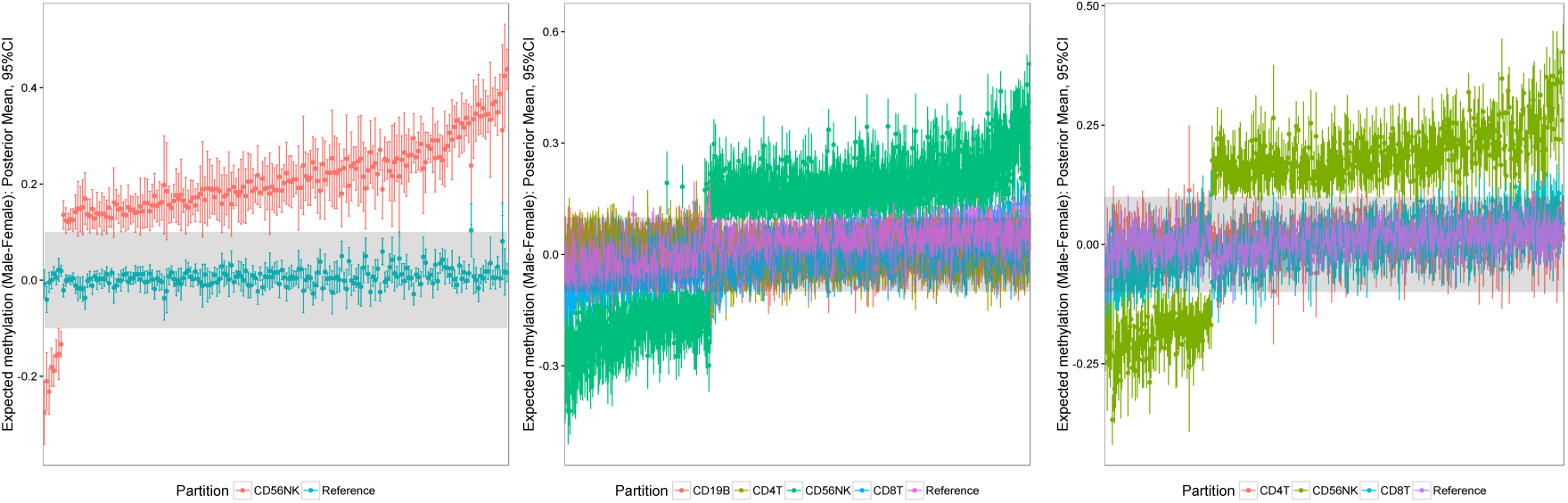
Posterior summaries of sex-specific differences in selected common marker panels defined by differences in CD56^+^ NK methylation only. The difference at each marker is summarised by the posterior mean and corresponding 95% CI. The shaded region corresponds to a difference of ±0.1. L-R: CD56^+^ NK, Lymphocyte-I, Lymphocyte-II. CI=Credible interval.

684 common Myeloid markers were associated with differences in CD14^+^ Monocytes or CD16^+^ Neutrophils only. Differences with respect to CD14^+^ Monocytes tended to be higher in females, compared with higher male methylation in CD16^+^ Neutrophils (Fig 6).

**Figure 6:**
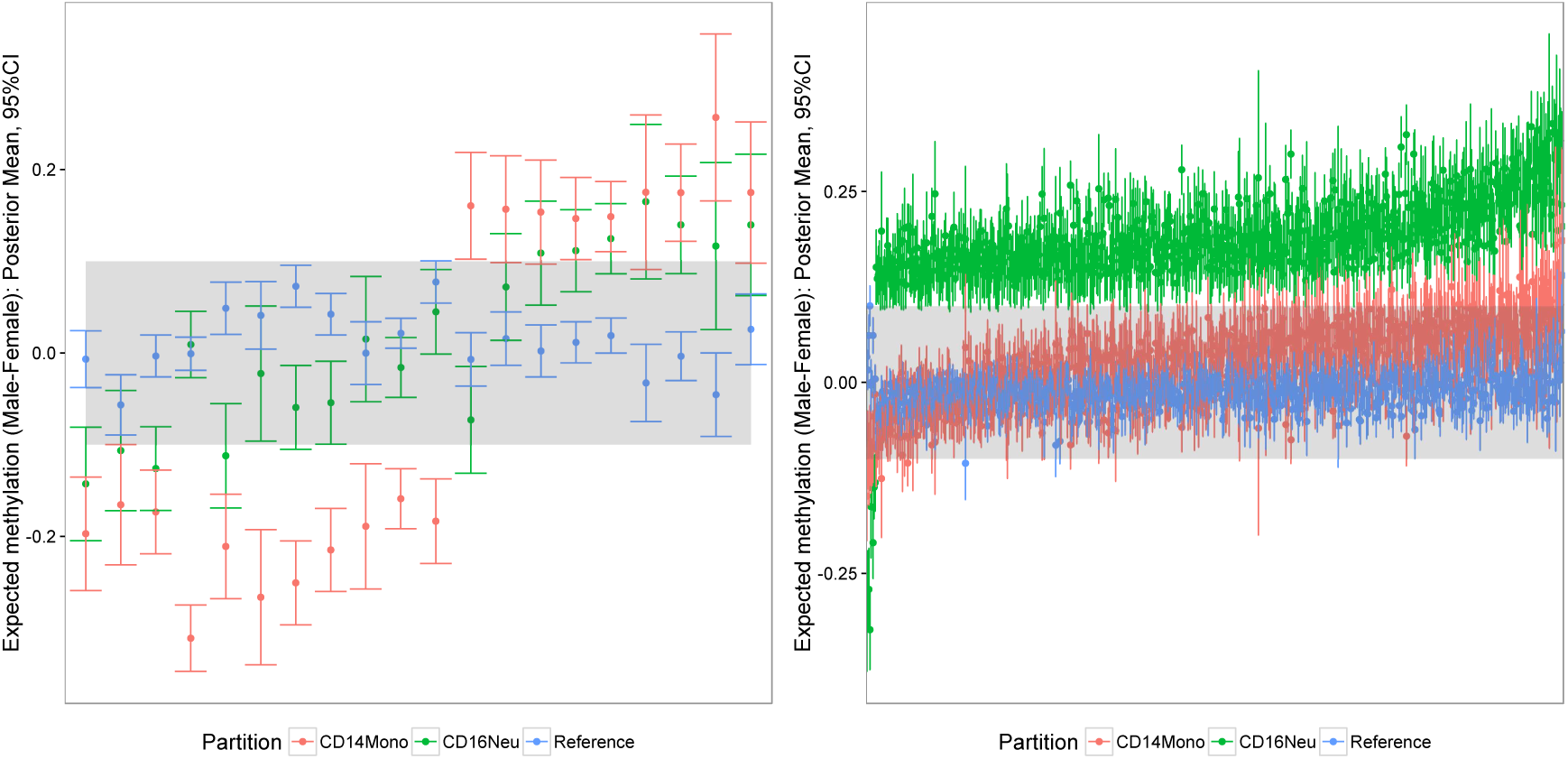
Posterior summaries of sex-specific differences in common Myeloid markers defined by differences in CD14^+^Monocytes (Left) or CD16^+^ Neutrophils (Right) only. The difference at each marker is summarised by the posterior mean and corresponding 95% CI. The shaded region corresponds to a methylation difference of ± 0.10.

## Discussion

This paper has proposed new statistical methodology for the discovery of cell-specific methylation profiles in whole blood, applying principles of Bayesian model selection. The characterisation of CpGs by differential methylation in one or more cell types builds significantly upon existing work that has been restricted to univariate analyses of differential methlyation by cell type. Sex-specific differences in both the prevalence of cell-specific profiles and methylation signal for select cell types were also demonstrated. For immune cell subtypes, validation of common markers using external cell-sorted samples produced favourable results. Enrichment analyses of common marker panels provided additional support for the proposed methodology, where it was demonstrated that the detection of cell-specific methylation at the individual CpG level was also biologically meaningful.

The incorporation of knowledge about hematopoietic lineage into model specification represents a semisupervised approach that reflects relevant cell biology. Consistent with other model selection strategies, our approach assumes that the defined set of candidate models is exhaustive and true patterns beyond this set are not of primary interest. In the event that the true methylation pattern is not commensurate with any of the candidate models specified, two outcomes are likely. In the first instance, corresponding probes areassigned to the saturated model and may be examined further as part of a larger marker panel. Alternatively, the vector of model probabilities for each CpG may be diffuse over multiple possibilities and therefore not be identified for any cell-specific pattern under the chosen criteria. The feasibility of unsupervised approaches to model selection, where the full list of methylation patterns is determined by the observed data is appealing, but their routine application in high dimensional settings is prohibitive. Furthermore, there is potential for patterns identified to be an artefact of random noise present in cell sorted samples as opposed to true biological signal. Integrated methods of model selection represent an avenue for future work, where new cell-specific patterns are proposed based on the combination of cell lineage and their prevalence in the observed methylome.

It is well documented that there are sex-specific differences in the proportions of circulating white blood cells [25, 26]. The application of the proposed methodology to female and male samples has highlighted the importance of accounting for sex effects in DNA methylation analyses. Greater numbers of CD19^+^B and T-cell dependent markers in females are consistent with previous findings and are possibly indicative of higher levels of cell activation [27]. The association between sex-specific differences in select CD4^^+^^T markers and the *CD40LG* gene have also been identified previously. Previous studies have pointed to alelle specific methylation for this gene [28, 29] where CD4^+^T hypomethylation is observed in healthy males compared with healthy women who carry one methylated and one hypomethylated alelle. One of our major findings was large differences of methylation between males and females in markers defined as CD56^+^NK specific. This is interesting when considered alongside the observation that males show an increase in circulatory NK cells compared to females [27], which adds further support for the accuracy of the approach. Additionally there is some evidence of sex-specific methylation differences in CD^+^56 NK, as well as CD^+^8 T-cells [30]. Under the proposed approach, we have provided a potential solution to accurately account for potential bias introduced by sex effects at the marker level.

The presence of cell-specific methylation CpG markers highlights the need to account for cellular composition prior to conducting Epigenome Wide Association Studies (EWAS), in whole blood. Methods for this purpose have been developed [31,32] based on the assembly of methylation ‘signatures’ from cell-sorted data which are then projected onto heterogeneous samples to predict cell type proportions. A comparison of common markers with the top 500 CpG probes identified by the cell mixture methodology of [31] revealed 75% concordance between panels (data not shown), of which the majority were associated with Lymphocyte-I/II and Myeloid maker panels. The inclusion of other marker panels in these algorithms may lead to further improvement in cell mixture estimation, in particular for immune cell subtypes that may be present in low proportions. Furthermore, the performance of these algorithms rely on consistent cell-type effects across cohorts [33]. Given the sex-specific methylation differences we have identified in this study, failure to account for sex effects may also impact upon the quality of cell mixture estimation and should therefore be given due consideration.

It is common practice in array-based methylation studies to exclude CpG sites which contain SNPs both within the probe and on the CpG site. While this is a valid approach to filtering before analysis, it will often lead to dramatic reduction of overall data. As a result, it is likely that sites of potential interest may be lost before any association can be made. By mapping hg38 annotated SNPs to all 450K CpG loci, we were able to ascertain the overall proportion of cell marker sites which have a SNP present; on average, across the common set of markers, this was approximately 30.7% of markers. In light of these results, we suggest that deconvolution studies and methods should account for SNP events at cell marker sites, noting the proportion that are present. For the filtering stage, we recommend that the overall rarity of the SNP variant be taken into account, for example, retaining CpGs which also have a ‘rare’ (MAF < 0.01) variant mapping. This approach is likely to be beneficial to the overall study design and outcome.

## Acknowledgments

We thank Peter Donnelly for his helpful feedback on the methodology developed in this paper. MB and NW were supported by a collaborative development grant from QUT. NW was further supported by the Australian Research Council (ARC) working under a Laureate Fellowship held by KM. DK is supported by an Australian Government Research Training Program (RTP) Scholarship, and an ARC Centre of Excellence for Mathematical and Statistical Frontiers (ACEMS) top-up scholarship. AF and RL are partially supported by Multiple Sclerosis Research Australia funding for bioinformatics.

## Supporting information

Supplementary files S1 and S2 are available from the GitHub respository https://github.com/nicolemwhite/BayesMS.

**S1 File** R code for Bayesian model selection algorithm. Core R functions required to prepare data and compute posterior model probabilities via the EM algorithm, across all listed candidate models.

**S2 File** List of common CpG markers associated with a sex-specific difference of ≥0.10. Illumina Human Methylation 450k annotation data are also included for each CpG marker.

**Table S1:** Percentage of markers associated with SNPs, for sex-specific and common markers by candidate model. The category ‘Unassigned’ refers to all CpG probes that were not assigned to any marker panel, based on a 5% Bayes’ FDR.

| Model | Female | Male | Common |
| --- | --- | --- | --- |
| CD19<sup>+</sup> B | 24.19 | 26.82 | 27.05 |
| CD14<sup>+</sup> Mono | 37.02 | 34.62 | 36.51 |
| CD4<sup>+</sup> T | 29.05 | 28.34 | 26.10 |
| CD8<sup>+</sup> T | 23.14 | 19.47 | 21.97 |
| CD16<sup>+</sup> Neu | 37.11 | 33.85 | 38.84 |
| CD56<sup>+</sup> NK | 33.70 | 36.82 | 36.02 |
| Pan T | 24.43 | 24.79 | 25.65 |
| Lymphocyte-I | 28.03 | 27.55 | 28.46 |
| Lymphocyte-II | 30.68 | 27.88 | 30.91 |
| Myeloid | 34.22 | 30.58 | 34.92 |
| All | 34.84 | 34.62 | 35.46 |
| Unassigned | 26.81 | 26.75 | 26.61 |

**Table S2:**
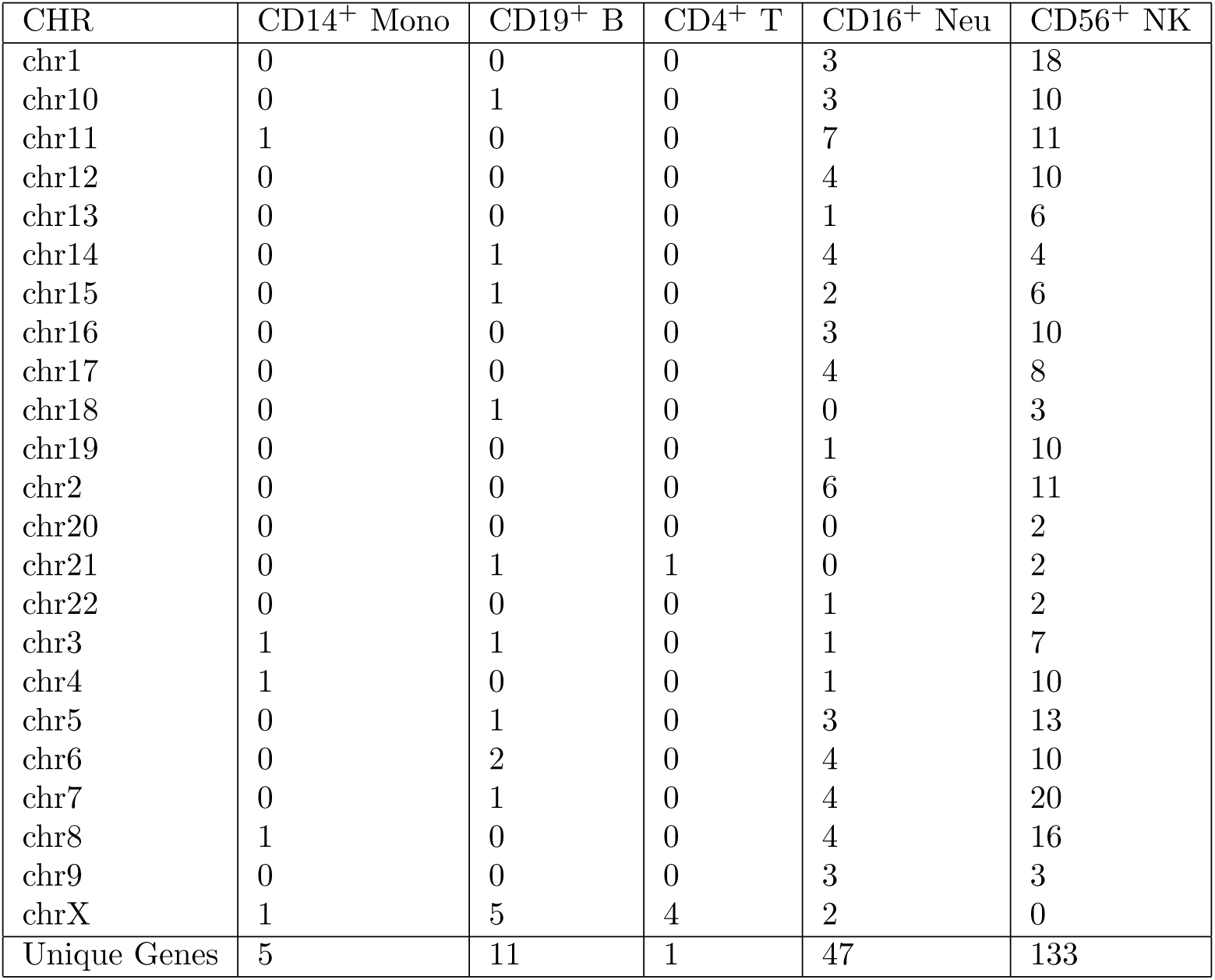
Distribution of sex-specific common markers over chromosomes, for single cell-dependent makers only. A sex-specific marker was declared if the posterior probability from Eq 3 was greater than 0.95 for at least one differentially methylated cell type.

| CHR | CD14<sup>+</sup> Mono | CD19<sup>+</sup> B | CD4<sup>+</sup> T | CD16<sup>+</sup> Neu | CD56<sup>+</sup> NK |
| --- | --- | --- | --- | --- | --- |
| chr1 | 0 | 0 | 0 | 3 | 18 |
| chr10 | 0 | 1 | 0 | 3 | 10 |
| chr11 | 1 | 0 | 0 | 7 | 11 |
| chr12 | 0 | 0 | 0 | 4 | 10 |
| chr13 | 0 | 0 | 0 | 1 | 6 |
| chr14 | 0 | 1 | 0 | 4 | 4 |
| chr15 | 0 | 1 | 0 | 2 | 6 |
| chr16 | 0 | 0 | 0 | 3 | 10 |
| chr17 | 0 | 0 | 0 | 4 | 8 |
| chr18 | 0 | 1 | 0 | 0 | 3 |
| chr19 | 0 | 0 | 0 | 1 | 10 |
| chr2 | 0 | 0 | 0 | 6 | 11 |
| chr20 | 0 | 0 | 0 | 0 | 2 |
| chr21 | 0 | 1 | 1 | 0 | 2 |
| chr22 | 0 | 0 | 0 | 1 | 2 |
| chr3 | 1 | 1 | 0 | 1 | 7 |
| chr4 | 1 | 0 | 0 | 1 | 10 |
| chr5 | 0 | 1 | 0 | 3 | 13 |
| chr6 | 0 | 2 | 0 | 4 | 10 |
| chr7 | 0 | 1 | 0 | 4 | 20 |
| chr8 | 1 | 0 | 0 | 4 | 16 |
| chr9 | 0 | 0 | 0 | 3 | 3 |
| chrX | 1 | 5 | 4 | 2 | 0 |
| Unique Genes | 5 | 11 | 1 | 47 | 133 |

**Figure S1:**
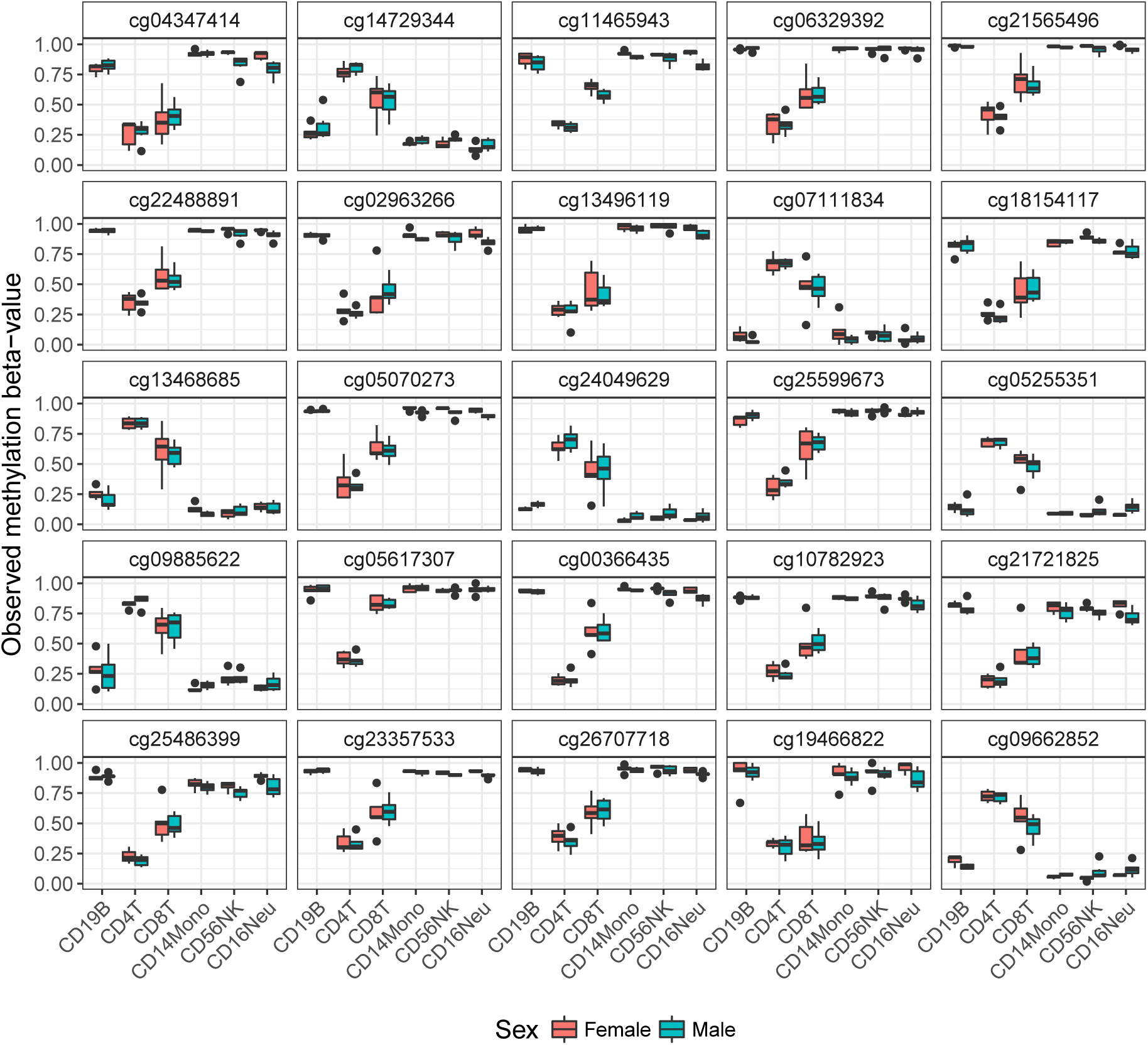
Distribution of observed methylation *β*–values by cell type and sex for selected common Pan T markers. Markers were identified as having high levels of differential methylation ( >0.5) in CD4^+^ T cells. Markers were identified if the corresponding posterior probability of differential methylation >0.5 exceeded 0.95.

**Figure S2:**
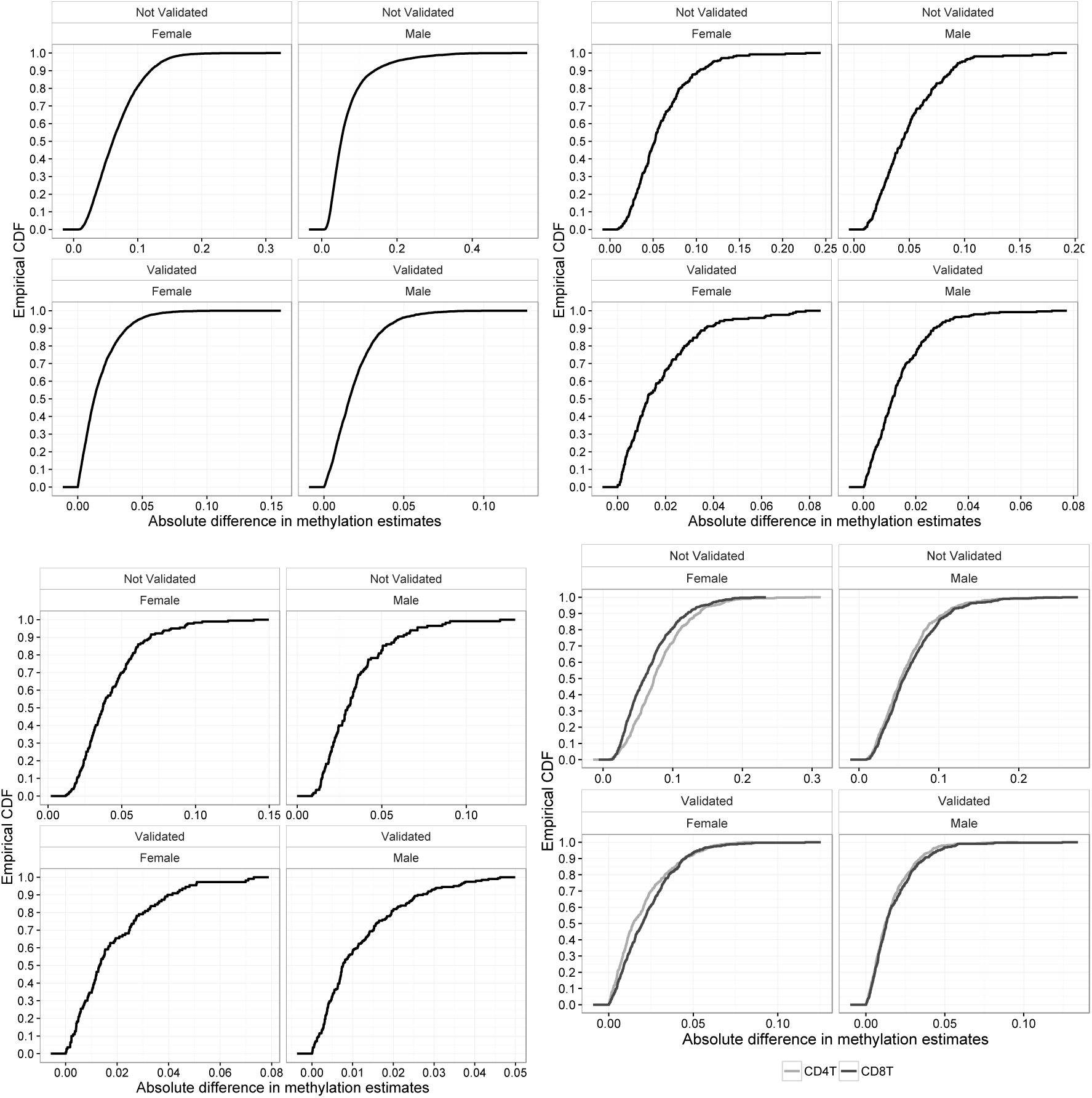
Validation analysis: Empirical cumulative distribution function (CDF) of the absolute difference between posterior and validation estimates by validation status. First row (L-R): CD19^+^B, CD4^+^T; Second row (L-R): CD8^+^T, Pan T.

## References

[1] Mary Muers. Gene expression: Disentangling DNA methylation. Nat Rev Genet, 14(8):519–519, jun 2013.

[2] Alexander Meissner, Tarjei S. Mikkelsen, Hongcang Gu, Marius Wernig, Jacob Hanna, Andrey Sivachenko, Xiaolan Zhang, Bradley E. Bernstein, Chad Nusbaum, David B. Jaffe, Andreas Gnirke, Rudolf Jaenisch, and Eric S. Lander. Genome-scale DNA methylation maps of pluripotent and differentiated cells. Nature, jul 2008.

[3] Tamar Hashimshony, Jianmin Zhang, Ilana Keshet, Michael Bustin, and Howard Cedar. The role of DNA methylation in setting up chromatin structure during development. Nature Genetics, 34(2):187–192, may 2003.

[4] S. Dedeurwaerder, M. Defrance, M. Bizet, E. Calonne, G. Bontempi, and F. Fuks. A comprehensive overview of Infinium HumanMethylation450 data processing. Briefings in bioinformatics, page bbt054, 2013.

[5] S. Dedeurwaerder, M. Defrance, E. Calonne, H. Denis, C. Sotiriou, and F. Fuks. Evaluation of the Infinium Methylation 450K technology. 2011.

[6] J. Sandoval, H. Heyn, S. Moran, J. Serra-Musach, M.A. Pujana, M. Bibikova, and M. Esteller. Validation of a DNA methylation microarray for 450,000 CpG sites in the human genome. Epigenetics, 6(6):692–702, 2011.

[7] V.K. Rakyan, H. Beyan, T.A. Down, M.I. Hawa, S. Maslau, D. Aden, A. Daunay, F. Busato, C.A. Mein, and B. et et al Manfras. Identification of type 1 diabetes–associated DNA methylation variable positions that precede disease diagnosis. PLoS Genet, 7(9):e1002300, 2011.

[8] Rodrigo Franco, Onard Schoneveld, Alexandros G. Georgakilas, and Mihalis I. Panayiotidis. Oxidative stress DNA methylation and carcinogenesis. Cancer Letters, 266(1):6–11, jul 2008.

[9] E.A Houseman, S. Kim, K.T. Kelsey, and J.K. Wiencke. DNA methylation in whole blood: uses and challenges. Current environmental health reports, 2(2):145–154, 2015.

[10] B.T. Adalsteinsson, H. Gudnason, T. Aspelund, T.B. Harris, L.J. Launer, G. Eiriksdottir, A.V. Smith, and V. Gudnason. Heterogeneity in white blood cells has potential to confound DNA methylation measurements.PloS one, 7(10):e46705, 2012.

[11] M.T. Bocker, I. Hellwig, A. Breiling, V. Eckstein, A.D. Ho, and F. Lyko. Genome-wide promoter DNA methylation dynamics of human hematopoietic progenitor cells during differentiation and aging. Blood, 117(19):e182–e189, 2011.

[12] John R Glossop, Nicola B Nixon, Richard D Emes, Kim E Haworth, Jon C Packham, Peter T Dawes, Anthony A Fryer, Derek L Mattey, and William E Farrell. Epigenome-wide profiling identifies significant differences in DNA methylation between matched-pairs of T- and B-lymphocytes from healthy individuals. Epigenetics, 8(11):1188–1197, nov 2013.

[13] L.E. Reinius, N. Acevedo, M. Joerink, G. Pershagen, S.E. Dahlén, D. Greco, C. Söderhäll, A. Scheynius, and J. Kere. Differential DNA methylation in purified human blood cells: implications for cell lineage and studies on disease susceptibility. PloS one, 7(7):e41361, 2012.

[14] Osman El-Maarri, Tim Becker, Judith Junen, Syed Saadi Manzoor, Amalia Diaz-Lacava, Rainer Schwaab, Thomas Wienker, and Johannes Oldenburg. Gender specific differences in levels of DNA methylation at selected loci from human total blood: a tendency toward higher methylation levels in males. Hum Genet, 122(5):505–514, sep 2007.

[15] Jingyu Liu, Marilee Morgan, Kent Hutchison, and Vince D Calhoun. A study of the influence of sex on genome wide methylation. PloS one, 5(4):e10028, 2010.

[16] F. Eckhardt, J. Lewin, R. Cortese, V. Rakyan, J. Attwood, M. Burger, T. Burton, J. and Cox, R. Davies, T. Down, et al. DNA methylation profiling of human chromosomes 6, 20 and 22. Nature genetics, 38(12):1378–1385, 2006.

[17] Nina S McCarthy, Phillip E Melton, Gemma Cadby, Seyhan Yazar, Maria Franchina, Eric K Moses, David A Mackey, and Alex W Hewitt. Meta-analysis of human methylation data for evidence of sex-specific autosomal patterns. BMC genomics, 15(1):1, 2014.

[18] Shimrat Mamrut, Nili Avidan, Elsebeth Staun-Ram, Elizabeta Ginzburg, Frederique Truffault, Sonia Berrih-Aknin, and Ariel Miller. Integrative analysis of methylome and transcriptome in human blood identifies extensive sex-and immune cell-specific differentially methylated regions. Epigenetics, 10(10):943–957, 2015.

[19] Arnold Zellner. On assessing prior distributions and Bayesian regression analysis with g-prior distributions. Bayesian inference and decision techniques: Essays in Honor of Bruno De Finetti, 6:233–243, 1986.

[20] F. Liang, R. Paulo, G. Molina, M.A. Clyde, and J.O. Berger. Mixtures of g priors for Bayesian variable selection. Journal of the American Statistical Association, 2012.

[21] M.A. Newton, A. Noueiry, D. Sarkar, and P. Ahlquist. Detecting differential gene expression with a semiparametric hierarchical mixture method. Biostatistics, 5(2):155–176, 2004.

[22] T Triche Jr. IlluminaHumanMethylation450k. db: Illumina Human Methylation 450k annotation data. R package version 2.0.9.

[23] J. Wang, D. Duncan, Z. Shi, and B. Zhang. WEB-based GEne SeT AnaLysis Toolkit (WebGestalt): update 2013. Nucleic Acids Research, 41(W1):W77–W83, may 2013.

[24] Y. Benjamini and Y. Hochberg. Controlling the false discovery rate: a practical and powerful approach to multiple testing. Journal of the Royal Statistical Society. Series B (Methodological), pages 289–300, 1995.

[25] Eleanor N. Fish. The X-files in immunity: sex-based differences predispose immune responses. Nature Reviews Immunology, 8(9):737–744, sep 2008.

[26] Sabra L. Klein and Katie L. Flanagan. Sex differences in immune responses.Nature Reviews Immunology, 16(10):626–638, aug 2016.

[27] M. Abdullah, P. Chai, M. Chong, E. Tohit, R. Ramasamy, C. Pei, and S. Vidyadaran. Gender effect on in vitro lymphocyte subset levels of healthy individuals.Cellular immunology, 272(2):214–219, 2012.

[28] Christian Schmidl, Maja Klug, Tina J Boeld, Reinhard Andreesen, Petra Hoffmann, Matthias Edinger, and Michael Rehli. Lineage-specific DNA methylation in T cells correlates with histone methylation and enhancer activity. Genome research, 19(7):1165–1174, 2009.

[29] Matlock Jeffries, Mikhail Dozmorov, Yuhong Tang, Joan T Merrill, Jonathan D Wren, and Amr H Sawalha. Genome-wide DNA methylation patterns in CD4+ T cells from patients with systemic lupus erythematosus. Epigenetics, 6(5):593–601, 2011.

[30] Masatoshi Inoshita, Shusuke Numata, Atsushi Tajima, Makoto Kinoshita, Hidehiro Umehara, Hidenaga Yamamori, Ryota Hashimoto, Issei Imoto, and Tetsuro Ohmori. Sex differences of leukocytes DNA methylation adjusted for estimated cellular proportions. Biology of Sex Differences, 6(1), jun 2015.

[31] Eugene Andres Houseman, William P Accomando, Devin C Koestler, Brock C Christensen, Carmen J Marsit, Heather H Nelson, John K Wiencke, and Karl T Kelsey. DNA methylation arrays as surrogate measures of cell mixture distribution. BMC bioinformatics, 13(1):1, 2012.

[32] D.C. Koestler, M.J. Jones, J. Usset, B.C. Christensen, R.A. Butler, M.S. Kobor, J.K. Wiencke, and K.T Kelsey. Improving cell mixture deconvolution by identifying optimal DNA methylation libraries (IDOL). BMC bioinformatics, 17(1):1, 2016.

[33] Paul Yousefi, Karen Huen, Hong Quach, Girish Motwani, Alan Hubbard, Brenda Eskenazi, and Nina Holland. Estimation of blood cellular heterogeneity in newborns and children for epigenome-wide association studies. Environmental and molecular mutagenesis, 56(9):751–758, 2015.

